# Protein Kinases in Phagocytosis: Promising Genetic Biomarkers for Cancer

**DOI:** 10.1101/2024.10.09.617495

**Authors:** Sadhika Arumilli, Hengrui Liu

## Abstract

Cancer is a complex disease characterized by genetic and molecular diversity, often involving dysregulation of critical cellular pathways. Recent advances in pan-cancer research have highlighted the importance of shared oncogenic mechanisms across different cancer types, providing new avenues for therapeutic exploration. Protein kinases, particularly those involved in phagocytosis, play pivotal roles in cellular homeostasis and immune response. This study systematically examines the genetic alterations and expression profiles of protein kinases associated with phagocytosis across various cancer types, using data from The Cancer Genome Atlas (TCGA) and other publicly available resources. We analyzed single nucleotide variations (SNVs), copy number variations (CNVs), methylation patterns, and mRNA expression to identify recurring alterations and their associations with survival outcomes. Our findings reveal that MET and MERTK are the most frequently mutated genes, with missense mutations dominating across cancers. CNV analysis shows significant correlations with survival in cancers like UCEC, KIRP, and KIRC, while methylation analysis indicates cancer-specific regulatory patterns affecting gene expression. Differential expression analysis highlights distinct cancer-type-specific expression profiles, with genes like MET and BTK displaying significant variation. Crosstalk pathway analysis further reveals the involvement of these kinases in key cancer-related pathways, such as epithelial-mesenchymal transition (EMT) and apoptosis. Drug sensitivity analysis identifies potential therapeutic targets, with gene expression correlating significantly with cancer cell line responsiveness to specific compounds. These findings underscore the importance of the phagocytotic kinome in cancer biology and suggest potential therapeutic strategies targeting protein kinases to enhance immune response and improve treatment outcomes.

## Introduction

The global cancer burden is expected to be 28.4 million cases in 2040, a 47% rise from 2020[1]. Traditionally, cancer research has focused on the tissue of origin, which can be limiting as it may fail to capture the underlying molecular and genetic diversity of tumors. The paradigm shift towards pan-cancer research has opened up new avenues of understanding. At the genetic and molecular levels, uncovering shared oncogenic pathways across different cancers has led to a deeper understanding of cancer biology and therapeutic targets.

Previous biomarker studies for cancer have been highly successful in identifying potential markers for cancer diagnosis and prognosis[2-7].

Protein kinase-mediated phagocytosis is increasingly recognized as a critical mechanism for maintaining cell homeostasis and innate immunity [8].Hence dysregulation of protein kinases in phagocytic cells can lead to impaired immune response which may contribute to the development and progression of cancer. CD47-SIRPa axis is such an emerging cancer therapy strategy9.Protein kinases belong to the phosphoryl-transferases superfamily of enzymes, which activate enzymes via phosphorylation[9-11]. The kinome of an organism is the total set of genes in the genome which encode all the protein kinases. The genome of protein kinases involved in phagocytosis (phagocytotic kinome) includes ABL1, AXL, BCR, BTK, CAMK1D, CSK, EIF2AK1, FGR, FYN, HCK, LIMK1, LYN, MERTK, MET, MST1R, PAK1, PRKCD, PRKCE, PRKCG, PTK2, SRC, SYK, and TYRO3.

In this study we set to identify recurring patterns of mutations and expression in the phagocytotic kinome of cancer cells and check for their statistical significance, thereby identifying pathways for future cancer therapy and targeted immunomodulation. Similar studies have been conducted to explore various gene sets across different cancer types[12-17]. We hope this work will contribute to further understanding in the role of Phagocytosis in cancer.

## Methods

### Data Acquisitions

Expression data, including clinical information, single nucleotide variants (SNVs), copy number variants (CNVs) and methylation data, were obtained from the Cancer Genome Atlas (TCGA) and the NCI Genomic Data Commons. Reverse phase protein array (RPPA) data were retrieved from the Cancer Proteome Atlas (TCPA). Immunotherapy response and survival data were retrieved from the TIDE (Tumor Immune Dysfunction and Exclusion) database. Gene-drug sensitivity data were collected from the Drug Sensitivity in Cancer (GDSC) database and the Cancer Therapeutics Response Portal (CTRP).

### Gene Alterations and Expression Analysis

SNV visualizations were generated with the Maftools package, while CNV data were processed with GISTIC2.0. Correlation between CNV and mRNA expression and between methylation levels and mRNA was assessed by Spearman correlation analysis. Differential analysis was performed to compare methylation patterns between tumor and normal samples, and individual variations in gene expression were also assessed. Gene Set Variation Analysis (GSVA) was applied to analyze mRNA expression data for the gene set using both Wilcoxon test and ANOVA t-test for statistical comparisons.

### Survival Analysis

The study examined mRNA expression, methylation, CNV, and clinical survival data. Tumor samples were divided into high and low groups based on median values or specific alterations, such as mutations and CNV classifications like amplification and deletion. Survival time and status were fitted using the R package survival, and the Cox Proportional-Hazards model was employed to calculate hazard ratios (HR) for each gene, indicating survival risk. Statistical significance of survival differences between groups was assessed using log-rank tests, and Kaplan-Meier survival analysis was conducted to further evaluate gene-specific survival impacts.

### Statistical Analysis

Statistical analyses were conducted using R software v4.0.3. The Spearman correlation test assessed correlations, and the Cox proportional hazards model calculated survival risk and hazard ratios (HR). Kaplan-Meier curves and log-rank tests evaluated prognostic values. T-tests or ANOVA were used for group comparisons, with a rank-sum test for two datasets unless specified otherwise. Significance was set at P<0.05. SNV.

### Pathway Analysis

Pathway activity values for 7,876 samples were calculated using reverse-phase protein array (RPPA) data from the TCPA database. The analysis covered ten cancer-associated pathways: hormone estrogen receptor (ER), hormone androgen receptor (AR), receptor tyrosine kinase (RTK), phosphatidylinositol-4,5-bisphosphate-3-kinase (PI3K)/protein kinase B (AKT), RAS/mitogen-activated protein kinase (MAPK), tuberous sclerosis complex (TSC)/mechanistic target of rapamycin (mTOR), epithelial-mesenchymal transition (EMT), cell cycle, and apoptosis pathways. Pathway scores were calculated by aggregating the relative concentrations of all positive regulatory proteins and subtracting those of negative regulators. To estimate Pathway Activity Scores (PAS), gene expression data were divided into high and low categories based on median values as in previous studies. PAS differences between these categories were assessed using Student’s t-test; p-value was adjusted using FDR, where FDR ≤ 0.05 was considered significant. A gene was found to exert an activating effect on a signaling pathway if PAS (Low expression of Gene A) < PAS (High expression of Gene A); otherwise, it was considered to exert an inhibitory effect.

### Immune Association Analysis

Immune cell infiltration levels within various cancers were analyzed using data from the TCGA database. The infiltrates of 24 immune cell types were evaluated using ImmuCellAI. Gene set variation analysis (GSVA) scores of the genes were used to visualize the data. The relationship between immune cell infiltration and gene expression was quantified using Spearman correlation analysis, with correlation coefficients indicating the strength of associations. P-values were adjusted using the false discovery rate (FDR).

### Drug sensitivity Analysis

Drugs were screened based on their CEPNA correlation with gene expression and drug sensitivity, using a stringent significance cutoff (p < 1e-5). The analysis employed GSCALite to calculate the area under the dose-response curve (AUC) values for drugs and the expression profiles of protein kinase genes involved in phagocytosis across various cancer cell lines. Drug sensitivity and gene expression data from the GDSC and CTRP databases were integrated for a comprehensive evaluation. Spearman correlation analysis was used to assess the relationship between gene expression and drug sensitivity.

## Results

### Single Nucleotide Variation Analysis

The analysis of single nucleotide variations (SNVs) in protein kinase genes associated with phagocytosis revealed that MET and MERTK were the most frequently mutated across different cancer types. MERTK exhibited the highest mutation frequency, with 51 SNV samples in 468 SKCM. Overall, UCEC had a higher SNV mutation rate across the genes, however, SKCM and COAD also presented significant mutation frequencies in several genes. (Figure 1A) The SNV landscape shows that the majority of the mutations were missense mutations, and the most common SNV classes were C>T and C>A. (Figure 1B) Additionally, the association between SNV of the genes and survival was analyzed. while some genes like TYRO3, LYN and PTK3 show significant correlations with survival in specific cancer types, most gene expressions do not exhibit strong associations with survival outcomes across the majority of cancer types analyzed. (Figure 1C)

**Figure 1:**
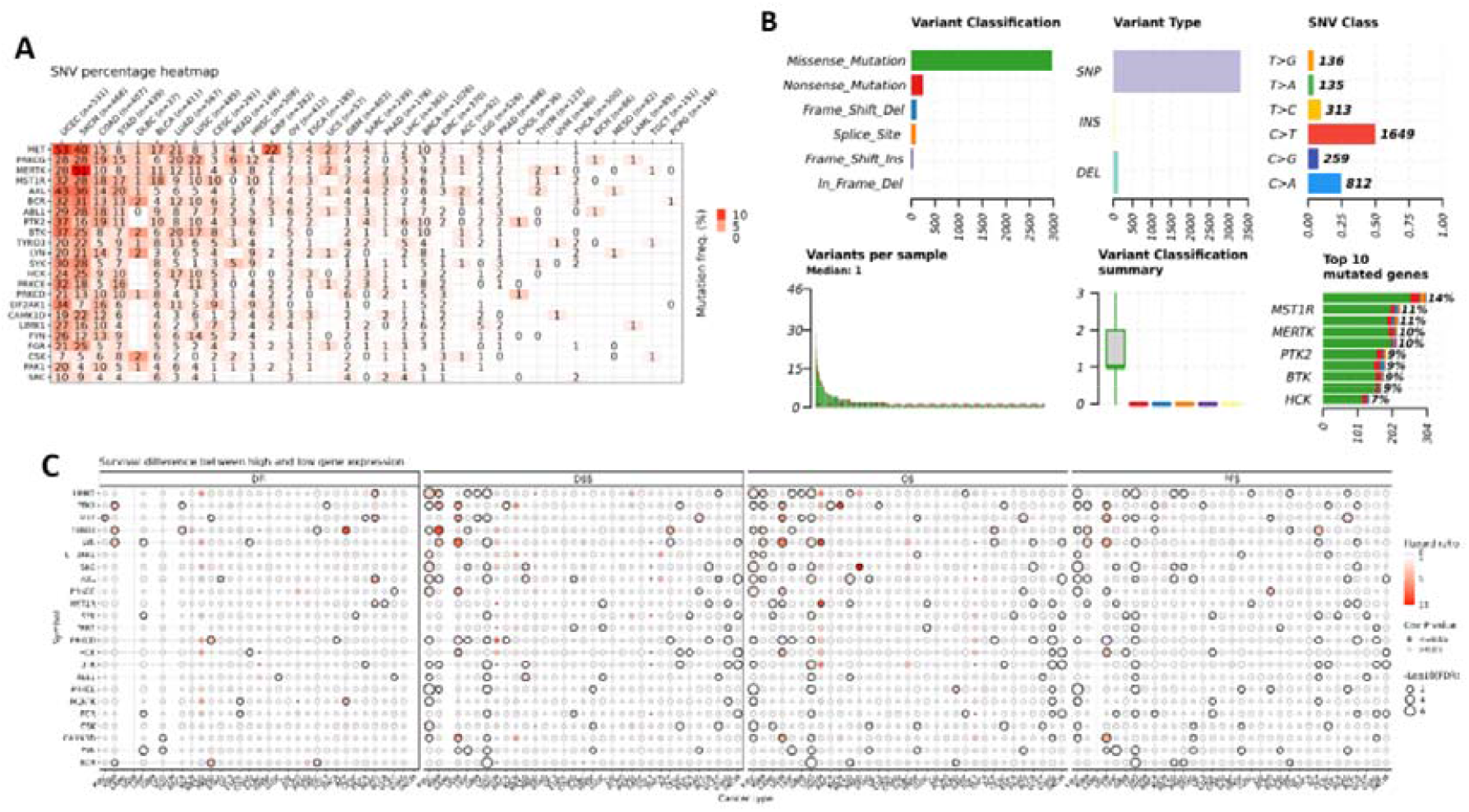
A – Heatmap of mutation frequency. The numbers indicate the count of samples exhibiting mutation frequencies within a specific cancer type; a value of 0 denotes the absence of mutation frequencies in that genomic region. The color represents the frequency of mutations per cancer type; B -An overview of SNV classes in protein kinase genes related to phagocytosis across cancers showing the count of each type of harmful mutation, the number of different variant types (SNP, INS, and DEL), the count of each SNV class, and the count of variants in each sample (represented by a bar, with the bar color corresponding to the variant classification) and a box plot shows the distribution of the count of each variant classification within the sample set (the box color matches the variant classification color) and the count and percentage of variants in the top 10 mutated genes; C – Correlation between SNV and survival in normal and tumor samples.

### Copy Number Variation Profile Analysis

Our analysis of the copy number variation (CNV) profile of the protein kinase genes associated with phagocytosis revealed diverse CNV patterns, with notable variations observed across different types of cancer. The distribution of CNV showed that heterozygous amplifications and deletions were predominant among the various CNV types (Figure 2A). We also plotted these CNV alterations to visualize their prevalence across cancer types (Figure 2B). Subsequently, we evaluated the correlation between CNV and gene expression. The cancers with the strongest correlations were BRCA, OV, LUAD, HNSC, and COAD; however, most cancers showed a positive correlation with expression (Figure 2C).

**Figure 2:**
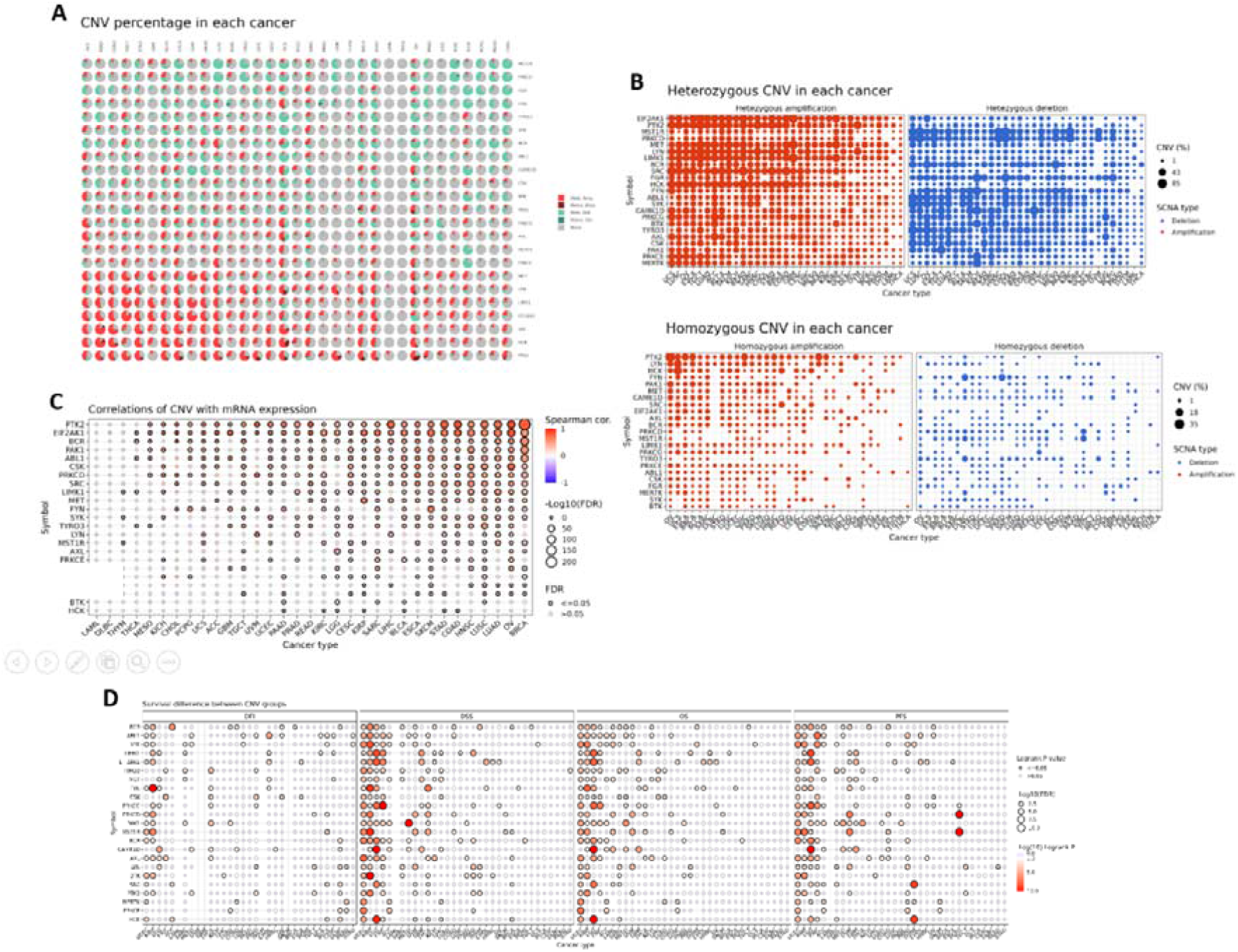
A - Pie charts of CNV distribution across cancers. Homo Amp = homozygous amplification; Homo Del = homozygous deletion; Hete Amp = heterozygous amplification; Hete Del = heterozygous deletion; B - Profile of heterozygous CNV and homozygous showing the percentage of amplification and deletion of the CNVs for each gene in each cancer; C – Correlation between CNV and “mRNA” expression; D-The survival difference between CNV groups.

Furthermore, we found significant associations between CNV and survival across various cancer types, especially in UCEC, KIRP, LGG, and KIRC (Figure 2D). This indicates that CNV levels in these cancers may be linked to differences in patient survival. Our analysis suggests a possible association between heterozygous CNV of protein kinase genes involved in phagocytosis and cancer.

### Methylation Analysis

The analysis of methylation differences between tumor and normal tissues revealed significant disparities. For instance, the genes MERTK and LYN show high methylation levels in BRCA while EIF2AK1 has low methylation levels in the same cancer type. The methylation differences are not uniform across all cancer types, indicating that each cancer type has a unique methylation profile for these genes (Figure 3A). The analysis of the correlation between methylation and mRNA expression profiles uncovered a consistent pattern across various cancer types, highlighting significant associations between methylation levels and expression which were mostly negative. However, there were certain genes like PTK2 and PRKCD in UVM which showed a strong positive association. (Figure 3B). In the survival profile analysis, only a few cancers demonstrated a relation between methylation levels and patient survival (Figure 3C). The analysis suggests that hypermethylation may lead to the down-regulation of specific genes, impacting their expression and potentially influencing patient outcomes in various cancer types.

**Figure 3:**
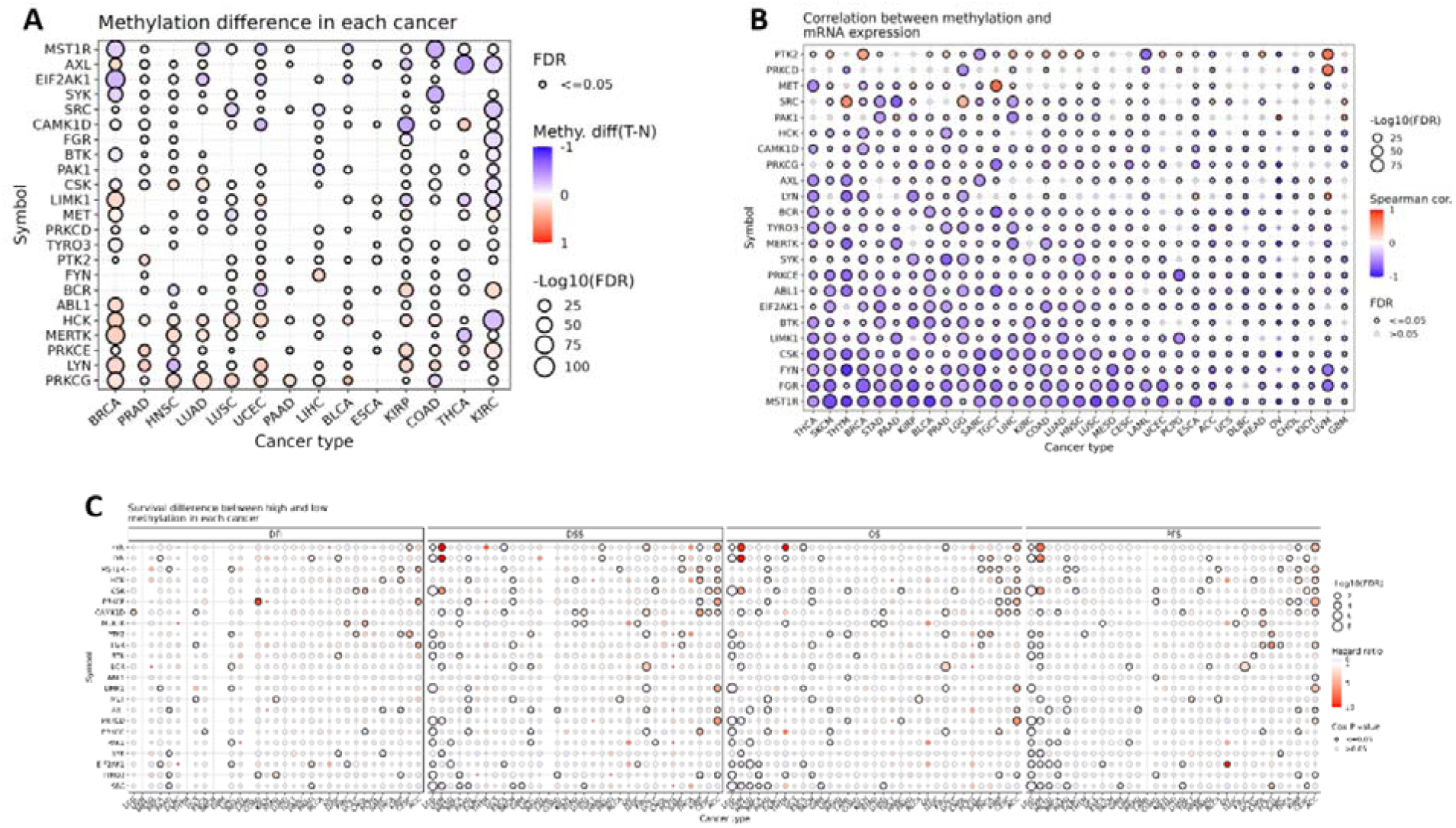
A - Difference in methylation between normal and tumor samples; B – The correlation between methylation and ‘mRNA’ expression; C – Difference in survival between samples with high and low methylation of the genes.

### mRNA Expression Analysis

The analysis of differential expression genes (DEGs) across various cancer types reveals some notable expression differences between cancer and non-cancer tissues. Using TCGA data, we examined the expression of protein kinase genes associated with phagocytosis in multiple cancer types (Figure 4A). In LUSC, genes like BTK, FGR, and PRKCE were significantly down-regulated. Conversely, in THCA, MET was up-regulated. Additionally, BRCA displayed a mix of up-regulated and down-regulated genes, with notable changes in EIF2AK1 and MET. Some cancer types, like KIRC and KIRP, exhibited both up-regulated and down-regulated genes. Overall, the expression profiles across cancer types indicate that while there are some consistently up-regulated or down-regulated genes in certain cancers, there are unique expression patterns that are specific to each cancer type. Nonetheless, for most cancer types, there were minimal notable variations in the expression of these genes between cancerous and normal tissues. In the analysis of gene expression subtype differences, BRCA and KIRC emerged as the most significant cancer types. However, STAD and LUAD also demonstrated notable significance (Figure 4B). The survival analysis revealed that KIRC and LGG were the top cancer types associated with these genes (Figure 4C). TYRO3 seems to have a higher survival risk when overexpressed in KIRP and ACC. It also shows that overexpression of LYN in UVM is associated with higher survival risk of patients.

**Figure 4:**
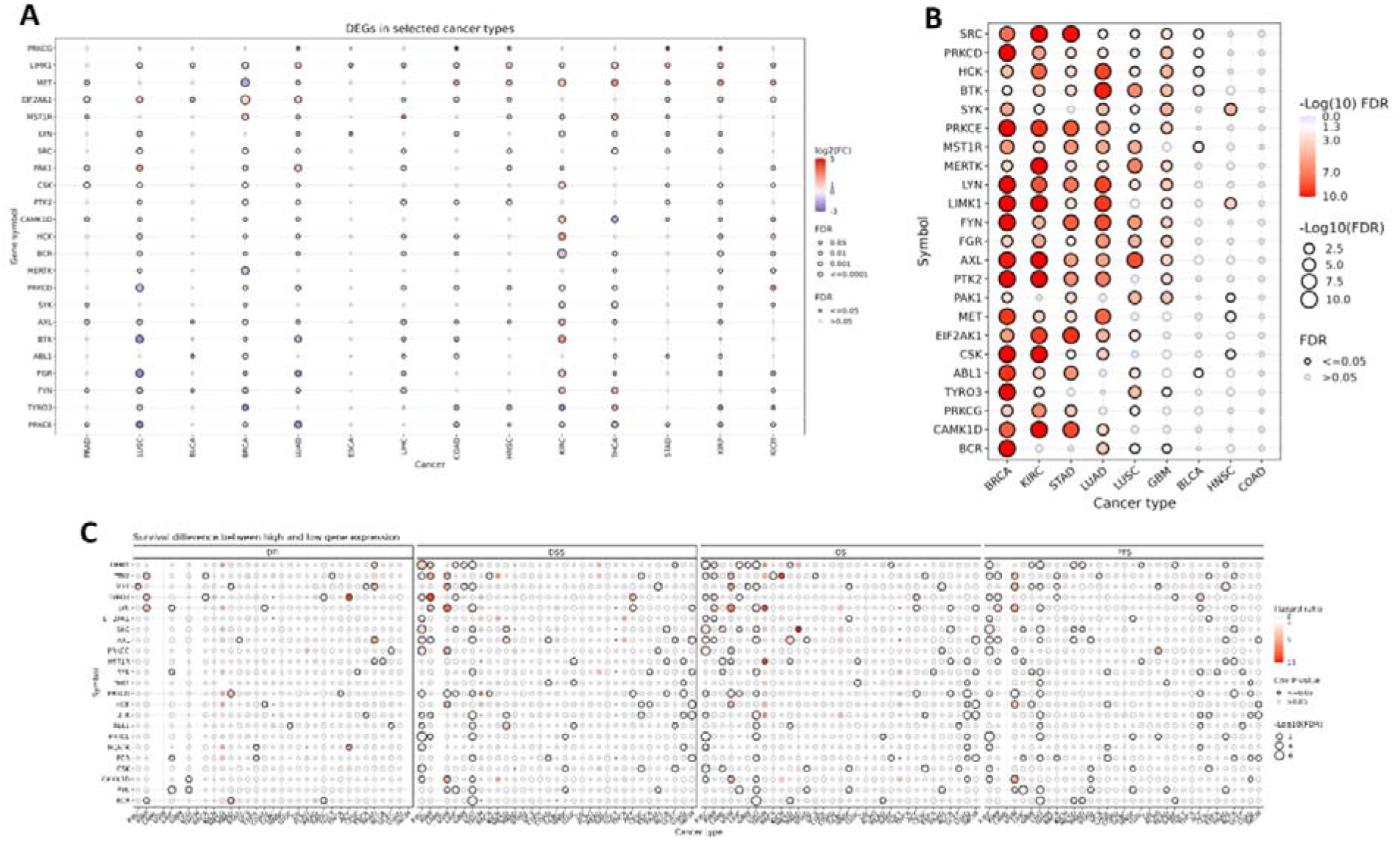
A - Comparism of mRNA expression levels between normal and tumour samples; B-Difference of expression levels between high and low subtypes; C – Difference of survival between high and low expression. The size of the dot represents the significance of the gene’s impact on survival across different cancer types, while the color indicates the hazard ratio. DFI: disease-free interval; DSS: disease-specific survival; OS: overall survival; PFS: progress-free survival.

### Crosstalk pathway activity analysis

To investigate the potential cross-talk of protein kinase genes involved in phagocytosis with other cancer-related pathways, we analyzed the data to determine the association of these genes with various biological processes. The analysis revealed that certain genes showed significant involvement in multiple pathways. Specifically, 44% of FYN, 41% of AXL and FGR, and 38% of HCK were associated with the activation of the epithelial-mesenchymal transition (EMT), suggesting a role in cancer migration and invasion (Figure 5). Additionally, 41% of AXL was linked to cell cycle inhibition, highlighting its potential role in controlling cell proliferation. In the context of apoptosis activation, genes such as CSK, LYN, LIMK1, and FGR showed associations, indicating their involvement in programmed cell death pathways. Furthermore, 38% of HCK was related to hormone estrogen receptor activation (ER_A), and both MET and LIMK1 showed associations with hormone androgen receptor inhibition (AR_I), suggesting a complex interplay between these protein kinases and hormone signaling pathways.

**Figure 5:**
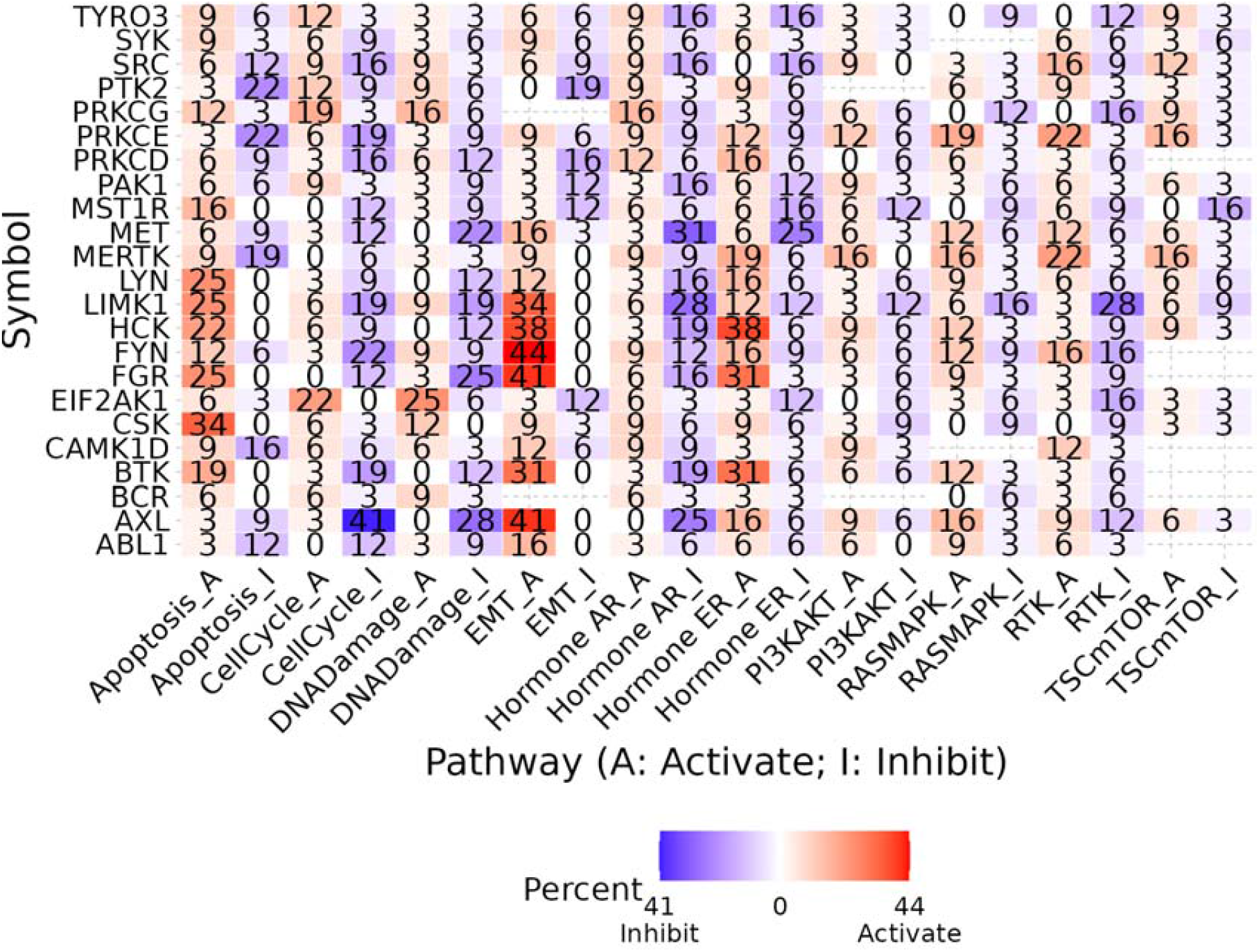
Correlation between expression and pathway activity of the genes.

### Drug Sensitivity Analysis

This study also investigated the relationship between protein kinase genes in phagocytosis and the tumor immune microenvironment. A gene set signature (GSVA score) was developed to evaluate the infiltration levels of different immune cells across various cancer types. The results indicated a significant correlation between the GSVA score and the infiltration levels of numerous immune cells. Particularly, the GSVA score showed a strong negative correlation with neutrophil cells and a positive correlation with macrophage cells (Figure 6A). Additionally, we examined the association between these protein kinase genes and drug sensitivity in cancer cell lines using data from the GDSC and CTRP databases. The results revealed that the protein kinase genes related to phagocytosis were significantly correlated with the sensitivity of cancer cells to multiple compounds (Figure 6B). These findings underscore the potential impact of protein kinase genes in phagocytosis on immune cell infiltration in cancers and suggest new avenues for targeted drug development.

**Figure 6:**
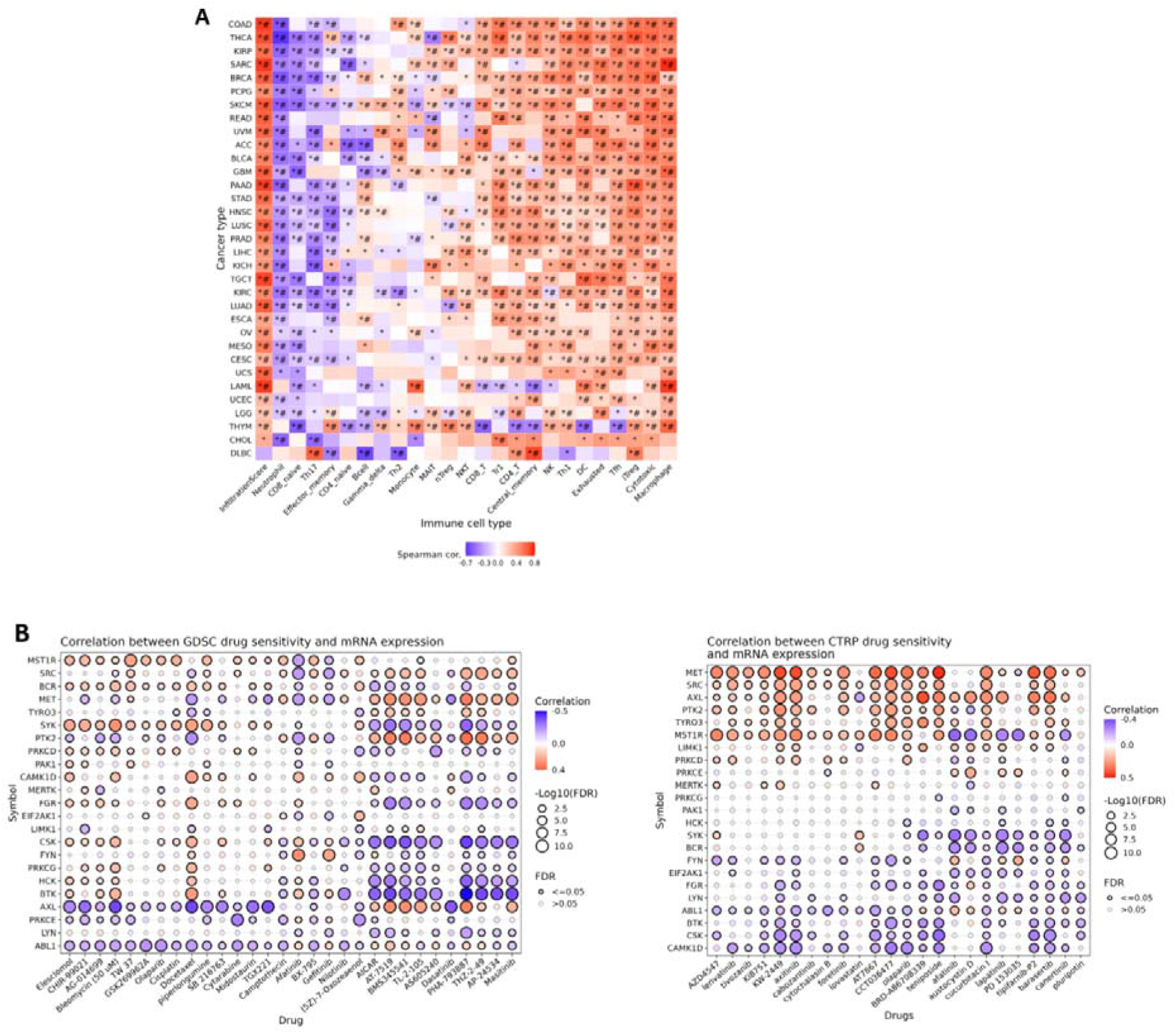
A - Correlation between GSVA scores of the genes and immune cell infiltration levels across cancer types, with statistical significance indicated by *p<0.05 and *#p<0.01. B - Correlation between the expression of the genes and the sensitivity of cancer cell lines to small molecules/drugs was analysed using GDSC and CTRP data.

## Discussion

The results of this study provide important insights into the genetic and molecular alterations of protein kinase genes involved in phagocytosis across various cancer types. These findings underscore the relevance of these genes not only in cancer progression but also in their potential as therapeutic targets.

The analysis of single nucleotide variations (SNVs) revealed that MET and MERTK were the most frequently mutated genes across multiple cancer types, with MET showing the highest mutation frequency in SKCM, while UCEC displayed an overall higher mutation rate. The presence of predominantly missense mutations, particularly in MET and MERTK, suggests these genes may be critical in the oncogenic processes of various cancers. These findings are consistent with previous studies that highlight MET’s role in cancer progression, where its mutation and overexpression drive tumor growth and metastasis. Furthermore, while genes such as TYRO3, LYN, and PTK3 exhibited significant correlations with survival in specific cancers, these associations were not universally observed across all cancer types, indicating a complex, context-dependent role of these kinases in cancer biology. Future studies should explore the mechanistic implications of these mutations to identify potential targets for therapeutic intervention.

Copy number variation (CNV) profiling revealed widespread heterozygous amplifications and deletions across cancer types, with significant survival associations observed particularly in UCEC, KIRP, and KIRC. The observed positive correlations between CNVs and gene expression in cancers like BRCA, OV, and LUAD suggest that CNVs play a critical role in modulating gene expression, which in turn may influence tumor behavior and patient outcomes. These findings align with previous reports that highlight CNVs as key drivers of cancer heterogeneity. The identification of significant CNV-survival associations further reinforces the clinical relevance of these alterations, suggesting that CNV profiling could serve as a valuable prognostic tool for personalized cancer treatment strategies.

Our methylation analysis highlighted cancer-specific patterns, with genes such as MERTK and LYN showing hypermethylation in BRCA and an inverse relationship between methylation levels and mRNA expression in most cancers. Interestingly, while hypermethylation was generally associated with gene silencing, a few exceptions, such as PTK2 in UVM, demonstrated a positive correlation between methylation and expression, suggesting alternative regulatory mechanisms at play. These findings imply that methylation may serve as a regulatory mechanism for protein kinase gene expression in cancer, with potential implications for early diagnosis and therapeutic targeting. The distinct methylation profiles observed across cancer types highlight the need for a cancer-type-specific approach when considering epigenetic therapies.

Differential expression analysis demonstrated significant cancer-specific variations in the expression of protein kinases associated with phagocytosis. While some genes such as MET were consistently upregulated in certain cancers like THCA, others, including BTK and FGR, were downregulated in LUSC. These results suggest that the phagocytotic kinome plays diverse roles in cancer biology, with specific genes either promoting or inhibiting tumor progression depending on the cancer type. The observed correlations between gene expression and survival, particularly for KIRC and LGG, further underscore the prognostic potential of these kinases, suggesting that targeting specific members of this gene family could offer new therapeutic avenues.

The crosstalk between the phagocytotic kinome and key cancer-related pathways, such as EMT and apoptosis, further emphasizes the importance of these kinases in tumor progression. The involvement of genes like FYN, AXL, and HCK in EMT highlights their potential role in promoting cancer cell migration and invasion. In contrast, the association of genes such as CSK and LYN with apoptosis underscores their role in modulating cell death pathways. These findings suggest that targeting the crosstalk between the phagocytotic kinome and these critical pathways could disrupt key processes in cancer development and progression, presenting new opportunities for therapeutic intervention. There are many other cancer related pathways such as MAPK[18], RAD-related pathways[19, 20], IGFBP-related pathway[21], etc, that should be studied in the future.

The observed correlations between protein kinase gene expression and drug sensitivity across multiple cancer cell lines indicate that these genes may influence cancer cell responsiveness to targeted therapies. In particular, the negative correlation between gene expression and sensitivity to certain compounds suggests that the overexpression of specific kinases may confer drug resistance. These findings offer valuable insights for the development of personalized cancer treatments, where targeting the phagocytotic kinome could enhance the efficacy of existing therapies or help overcome drug resistance. Bacteria such as Helicobacter pylori have been reported to potentially impact cancer drug sensitivity[22]. Further efforts are needed to better understand these mechanisms.

While this study provides a comprehensive overview of the genetic and molecular alterations of protein kinases involved in phagocytosis across cancers, several limitations must be acknowledged. First, the reliance on publicly available datasets such as TCGA may introduce biases related to sample composition and data quality[23]. Additionally, while this study identified significant associations between genetic alterations and survival, further experimental validation is required to establish causal relationships.

Future research should focus on the functional characterization of these protein kinase genes in cancer models to better understand their roles in tumor progression and immune modulation. Moreover, Single-cell analysis should be conducted in the future to better understand immune cells in tissue, as previous studies have successfully done[24]. Integrating multi-omic data, including proteomics and metabolomics[25], could provide a more comprehensive understanding of the phagocytotic kinome’s role in cancer. Finally, experimental validation of the function of these genes should be performed, similar to previous studies on viability or cell migration[26-29].

Last, traditional Chinese medicine (TCM) has been widely reported for its therapeutic potential in various human diseases[30-35]. Emerging studies suggest that TCM might also be beneficial for cancer treatment[36, 37]. However, it remains unclear whether protein kinases involved in phagocytosis can be regulated by TCM. This area warrants further investigation in future studies.

## Declarations

### Author Contributions

Sadhika Arumilli conducted all analyses and drafted the manuscript. Hengrui Liu designed the study, supervised the project, provided guidance to Sadhika Arumilli, and edited the manuscript.

## Acknowledgments

This study was conducted during the Lumiere project and received support from the Lumiere Company. The author thanks the support of Weifen Chen, Zongxiong Liu, Yaqi Yang, and Bryan Liu.

## Availability of data and materials

The source of the raw data was provided in the paper and the raw analysis data of this study are provided by the corresponding author with a reasonable request.

## Ethical approval

Not applicable.

